# Phylogenomics of *Mycobacterium africanum* reveals a new lineage and a complex evolutionary history

**DOI:** 10.1101/2020.06.10.141788

**Authors:** Mireia Coscolla, Daniela Brites, Fabrizio Menardo, Chloe Loiseau, Sonia Borrell, Isaac Darko Otchere, Adwoa Asante-Poku, Prince Asare, Leonor Sánchez-Busó, Florian Gehre, C. N’Dira Sanoussi, Martin Antonio, Affolabi Dissou, Paula Ruiz-Rodriguez, Janet Fyfe, Erik C. Böttger, Patrick Becket, Stefan Niemann, Abraham S. Alabi, Martin P. Grobusch, Robin Kobbe, Julian Parkhill, Christian Beisel, Lukas Fenner, Conor J. Meehan, Simon R Harris, Bouke C. De Jong, Dorothy Yeboah-Manu, Sebastien Gagneux

## Abstract

Human tuberculosis is caused by members of the *Mycobacterium tuberculosis* Complex (MTBC). The MTBC comprises several human-adapted lineages known as *M. tuberculosis* sensu stricto as well as two lineages (L5 and L6) traditionally referred to as *M. africanum*. Strains of L5 and L6 are largely limited to West Africa for reasons unknown, and little is known on their genomic diversity, phylogeography and evolution. Here, we analyzed the genomes of 365 L5 and 326 L6 strains, plus five related genomes that had not been classified into any of the known MTBC lineages, isolated from patients from 21 African countries.

Our population genomic and phylogeographical analyses show that the unclassified genomes belonged to a new group that we propose to name MTBC Lineage 9 (L9). While the most likely ancestral distribution of L9 was predicted to be East Africa, the most likely ancestral distribution for both L5 and L6 was the Eastern part of West Africa. Moreover, we found important differences between L5 and L6 strains with respect to their phylogeographical substructure, genetic diversity and association with drug resistance. In conclusion, our study sheds new light onto the genomic diversity and evolutionary history of *M. africanum,* and highlights the need to consider the particularities of each MTBC lineage for understanding the ecology and epidemiology of tuberculosis in Africa and globally.

## Introduction

Tuberculosis (TB) causes more human deaths than any other infectious disease, and it is among the top ten causes of death worldwide (*1*). Among the 30 high TB burden countries, half are in Sub-Saharan Africa (*1*). Africa also comprises the highest number of countries with the highest TB mortality (*1*). TB in humans and animals is caused by the *Mycobacterium tuberculosis* Complex (MTBC) (*2*), which includes different lineages, some referred to as *Mycobacterium tuberculosis* sensu stricto (Lineage 1 to Lineage 4 and Lineage 7) and others as *Mycobacterium africanum* (Lineage 5 and Lineage 6), a recently discovered Lineage 8 (*3*), as well as different animal-associated ecotypes such as *M. bovis*, *M. pinnipedii*, or *M. microti* among others (*4*, *5*). Among the human-associated MTBC lineages, some are geographically widespread and others more restricted (*6*). The latter is particularly the case for Lineage (L) 7 that is limited to the Horn of Africa (*7*, *8*), and L5 and L6 that are mainly found in West Africa (*9*). L5 and L6 differ substantially from the other lineages of the MTBC with respect to metabolism and in vitro growth (*10*, *11*). Several mutations in different genes of the electron transport chain and central carbon metabolic pathway can explain metabolic differences between L5 and L6 and the other lineages (*12*). L5 and L6 are also less virulent than other lineages in animal models, and appear to transmit less efficiently in clinical settings (*13*, *14*). Even though L5 and L6 are mostly restricted to West-Africa, they show a prevalence of up to 50% among smear-positive TB cases in some West African countries (*15*–*18*). Hence, L5 and L6 contribute significantly to the overall burden of TB across sub-Saharan Africa. Compared to the other MTBC lineages, relatively little is known with regard to the ecology and evolution of L5 and L6 (*5*, *19*). Two studies have found L5 to be associated with Ewe ethnicity in Ghana (*20*, *21*), supporting the notion that this lineage might be locally adapted to this particular human population (*22*). Several epidemiological associations suggest that L6 might be attenuated for developing disease as compared to other lineages (reviewed in (*9*)). For example, L6 has been associated with slower progression from infection to disease in The Gambia (*19*). Other studies have linked L6 with HIV co-infection in TB patients from The Gambia and Ghana (*19*, *21*), although other studies in Ghana and Mali have not seen such an association (*23*, *24*). Human TB caused by *M. bovis* compared to *M. tuberculosis* has also been associated with HIV (*25*) and higher levels of immunosuppression as CD4 T cell counts ≤200 cells/μL (*26*), leading to the suggestion that L6 might be an opportunistic pathogen, similar to *M. bovis* in humans (*27*). L5 and L6 also differ in various molecular features relevant for patient diagnosis, such as a non-synonymous mutation in the MPT64 antigen (*28*) and reduced T cell response to ESAT6 (*29*), leading to reduced detection by interferon gamma release assays of L5 and L6 in clinical samples (*28*, *30*). To shed more light on the phylogeography, evolutionary history and population genetic characteristics of *M. africanum*, we analysed the largest set of whole genome data for L5 and L6 generated to date.

## Results

### New MTBC Lineage: Lineage 9

We analysed a total of 696 *M. africanum* genomes. These included 365 L5 and 326 L6 genomes, as well as five related genomes that could not be classified into any of the known human- or animal-associated MTBC lineages (*4*, *31*). Out of these 696 genomes, 662 (95%) came from patient isolates originating in one of 21 countries of Sub-Saharan Africa. Another 34 (5%) strains were isolated outside Africa from patients with an origin other than Africa, or unknown (Table S1). To have a representative dataset and avoid overrepresentation of clustered strains, we removed 272 isolates that were redundant, whiles keeping the maximum phylogenetic diversity (>95% of the tree length) (*32*). The resulting non-redundant dataset comprised 424 genomes and showed a similar country distribution compared to the original dataset (Fig. S1).

We first focused our analysis on the five genomes that could not be classified into any of the known MTBC lineages. To explore the evolutionary relationship of these five genomes in the context of *M. africanum* diversity, we constructed the phylogeny of the 424 *M. africanum* genomes plus a reference dataset of animal associated MTBC genomes we published previously (*4*). The resulting phylogeny (Fig. 1) corroborated the separation of L5 and L6, and the localization of L6 in a monophyletic clade together with the animal-associated lineages, as previously described (*4*).

**Fig. 1.**
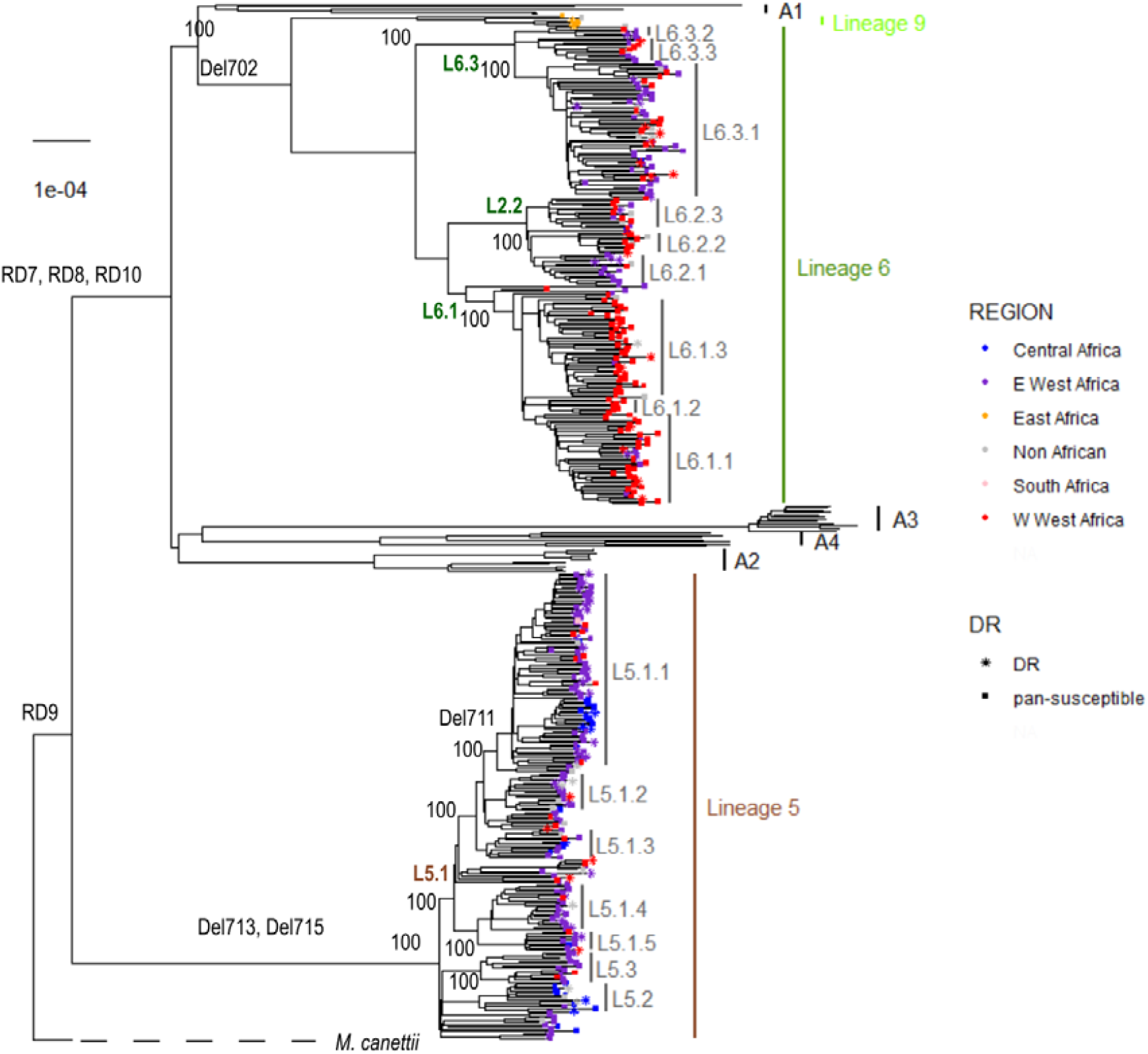
Maximum likelihood phylogeny of 424 *M. africanum* genomes analysed together with reference animal associated genomes. Support bootstrap values are indicated at the nodes. Nodes are coloured according to country or origin, and shape of the node indicates susceptible or drug resistance based on absence or presence at least one of the drug resistance mutations indicated in Table S8.

To further explore the phylogenetic position of these five genomes we constructed a genomic phylogeny with 248 reference genomes (*3*) including all eight human associated lineages and four animal associated lineages (Fig. 2). The five unclassified genomes appeared as a sister clade of L6, branching between L6 and the animal clade A1 (Fig. 2). The geographical origin of the five genomes differed from all other *M. africanum* genomes included in our analysis, as they were the only ones isolated from patients originating in East Africa (one from Djibouti, three from Somalia and one isolated in Europe but patient origin was unknown). By contrast, all L5 and L6 genomes came from patients originating in either West Africa (354 genomes) or Central Africa (37 genomes), except for one isolated in South Africa (Fig. 1) and 28 isolated outside Africa and from unknown origin.

**Fig. 2.**
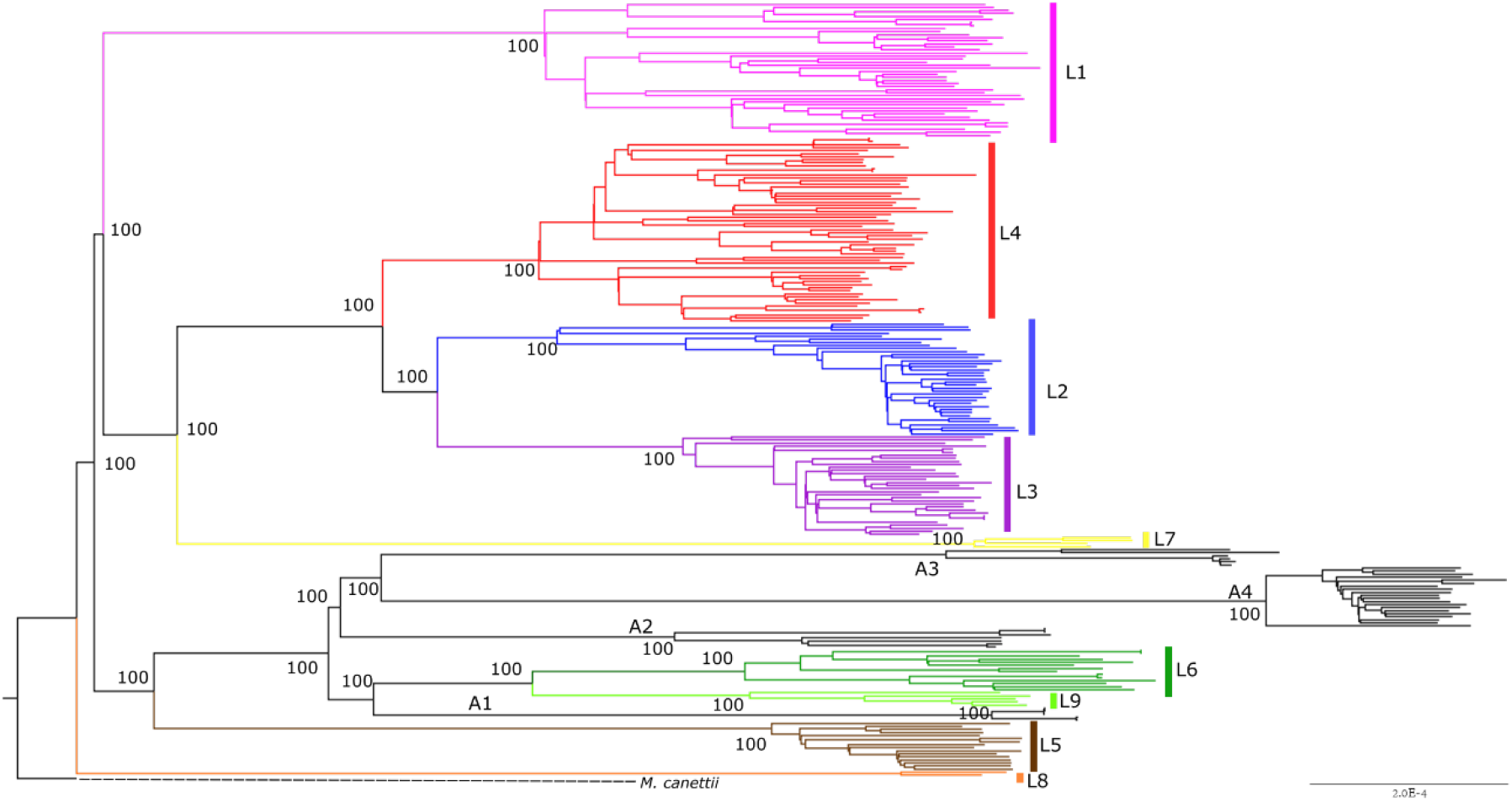
Maximum likelihood phylogeny of five unclassified genomes analysed together with reference dataset of MTBC genomes. The five unclassified genomes are coloured in light green and tagged as “L9”. Animal associated lineages A1 to A4 are indicated and coloured in black. Support bootstrap values are indicated at the deepest nodes.

The five unclassified genomes showed the following in silico inferred spoligotype: 772000007775671 (nnnnnnonoooooooooooooooonnnnnnnnnnonnnonnnn) in the genome from Djibouti, 772700000003671 (nnnnnnononnnoooooooooooooooooooooonnnnonnnn) in all three Somalian genomes, and a very similar pattern 772600000003631 (nnnnnnononnooooooooooooooooooooooonnnnoonnn) in the genome from Europe, for which the patient origin was unknown. We searched for these three spoligotypes in the international genotyping database SITVIT2, which includes 9,658 different spoligotypes from 103,856 strains isolated in 131 countries (*33*). Spoligotype 772600000003631 was not found among the 103,856 strains included in the database, and the other two spoligotypes can be considered extremely rare because they have been found only in three strains in the database: 772000007775671 in a strain isolated in France, and 772700000003671 in two strains isolated in the Netherlands, although patient’s origin is unknown.

The five unclassified genomes showed a mean distance of 1,191 SNPs to L6 genomes, 1,632 SNPs to L5 genomes, and 1,491 SNPs to the animal-associated MTBC genomes. Those distances were higher than the corresponding intra-lineage differences: 342 (standard deviation (SD) 3.65) within L5, 542 (SD 9.19) within L6, and 332.4 (SD 14.48) within the unclassified genomes. So, even when correcting for the diversity within each lineage, we still found that the five unclassified genomes were separated from the other lineages by 1,294, 582 and 654 SNPs of net distance to L5, L6 and the animal-associated lineages, respectively. Given the different geographical distribution and the substantial genetic separation, we classified these five genomes into a new MTBC lineage that we propose to call MTBC Lineage 9 (L9). The average intra-lineage diversity among these five L9 strains was 332 SNPs (SD=13). The maximum diversity within L9 was 514 SNPs between strain G00075 and strain G00074, with the smallest distance being 99 SNPs between strain G04304 and strain G00075.

We looked for deleted regions in the L9 genomes that could be used as phylogenetic markers, as was done for other MTBC lineages in the past (*34*, *35*) (*6*). We identified one region deleted in all L9 genomes that spanned from Rv1762c to Rv1765. However, this region is not a robust phylogenetic marker because partially overlapping deletions can be found in other lineages. Specifically, Rv1762c is deleted in genomes from one of the animal associated lineages, Lineage A3, which includes the strain previously known as *M. orygis,* and the region between Rv1763c and Rv1765 is deleted in L6 genomes. Hence instead, we report a list of SNPs that can be used as phylogenetic markers for L9 (Table S2) given that they appear in all five L9 genomes and are absent from genomes from other lineages (*32*).

Given the low number of L9 genomes, we focused the remaining of our analysis on *M. africanum* L5 and L6.

### Sublineages within L5 and L6

Our extended genomic analysis of L5 and L6 confirmed the deletions of the previously described regions of difference (RDs), including RD7, RD8, RD9 and RD10 (*34*, *35*), and RD713 and RD715 (*6*) as indicated in the phylogeny (Fig. 1). However, the deletion of RD711 could not be confirmed as a L5 marker as proposed previously (*6*), as it was only deleted in a subset of L5 genomes as reported recently (*36*). We found RD711-deleted genomes to form a monophyletic clade within L5; named L5.1.1 considering previous nomenclature as proposed in Ates et al. (*36*). In contrast, RD702 was found to be deleted in all L6 strains as shown previously (*6*), as well as in the newly defined L9 strains (Fig. 1).

Our phylogeny revealed a different topology for L5 as compared to L6. Specifically, the L5 phylogeny showed little structure. Nevertheless, we managed to subdivide L5 into three main sublineages that were well differentiated and highly supported by bootstrap values >90, and named them consistent with previous nomenclature (*36*) as L5.1, L5.2 and L5.3. Due to the high genomic diversity within L5.1, this group was further subdivided into five subgroups (Fig. 1), leading to a total of seven sublineages in this first and second level of subdivision. Sublineage classification was only partially corroborated by the results of the PCA performed on whole genome SNPs (Fig. 3A), where these sublineages were not clearly separated. By contrast, L6 showed a more differentiated tree structure with three clearly differentiated monophyletic sublineages (L6.1, L6.2 and L6.3) at the first level that could be further subdivided into a second level subdivision with at least three other subgroups each (Fig. 1), resulting in a total of nine sublineages. The first level of subdivision was strong for L6, where L6.1, L6.2 and L6.3 were clearly separated using PCA (Fig. 3B). However, sublineages at the second level of subdivision were not that clearly separated (Fig. 3B). To explore the robustness of the classification beyond PCA, we estimated genetic differentiation for each of these sublineages using the fixation index (FST) based on Wright’s F-statistic (*37*) as measure of population differentiation due to genetic structure. We conducted a hierarchical analysis comparing the population structure at the two levels of subdivision: one level with the three main groups for both L5 and L6, and a second level with all seven and nine sublineages of L5 and L6, respectively. The L5 population structure showed the highest differentiation within all seven sublineages, where the highest population differentiation index Fst=0.48 (p-value<0.000001), and the lowest population differentiation index was found between the three main sublineages at the first level of subdivision (Fst=0.14, p-value=0.04915). Similarly, Fst between all seven L5 sublineages showed moderate differentiation with pairwise Fst values between 0.3 and 0.5 (Table S3) and net pairwise differences between 76 and 206 SNPs (Table S4). Conversely, for L6, the higher differentiation was at the first level of subdivision, that is between the main sublineages (L6.1, L6.2, L6.3, with 47% of the variation, Fst=0.47, p=0.0035), mirroring the PCA results. Even differentiation at the second level of subdivision, that is between all nine sublineages of L6, showed more structure than for L5, with Fst values ranging between 0.25 and 0.75 (Table S5), and net pairwise differences of between 73 and 493 SNPs (Table S6). A list of SNPs found exclusively in each of the L5 and L6 sublineages is shown in Table S7.

**Fig. 3.**
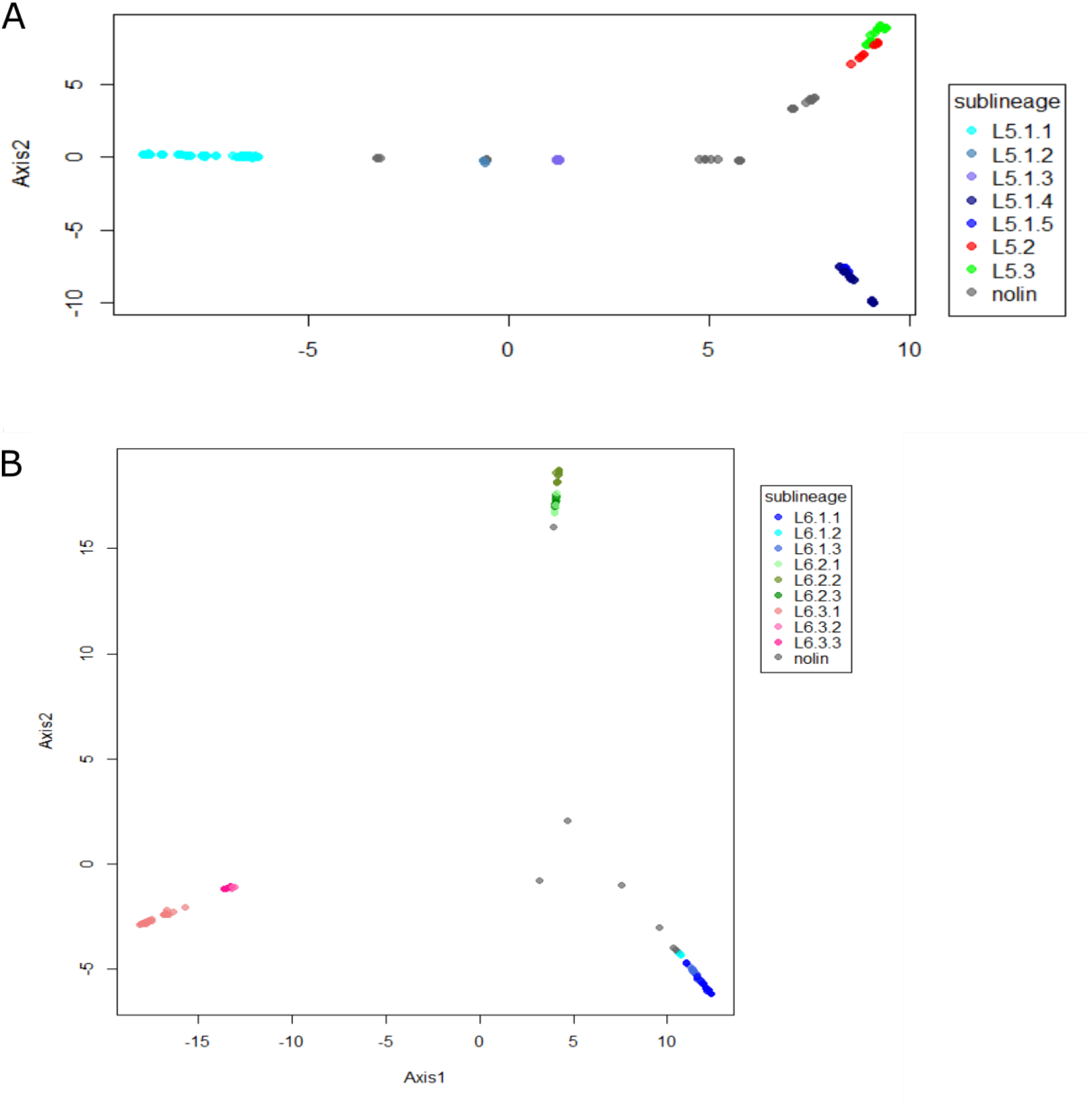
Principal Component Analysis (PCA) based on genomic variable SNPs. The PCA was conducted separately for L5 (A) and L6 (B). Colours indicate different sublineages and grey indicates genomes with no sublineage assigned “nolin”.

### Phylogeography

To explore the phylogeographical structure of L5, L6, and L9, we mapped the geographical origin of the genomes onto the phylogenetic tree as a coloured point at the end of each branch (Fig. 1). We grouped the different countries represented in the dataset into five regions in Africa: East, South, Central, and the Western part of West Africa (^W^West Africa) and the Eastern part of West Africa (^E^West Africa). We observed that most sublineages showed a characteristic geographical association at the regional level. At least five sublineages within L6 (all three L6.1 and two L6.2) showed a majority of genomes originating in ^w^West Africa, mostly The Gambia. By contrast, a few scattered L6 genomes, one sublineages within L6.2 and all three L6.3 genomes came from ^E^West Africa, mostly Ghana. Only a few L6 strains were found in Central Africa (N=2) or outside Africa (N=15). L5 showed a different phylogeographical structure with most genomes originating in ^E^West Africa (mostly Ghana) and two groups (L5.2 and one sublineage within L5.1.1) in Central Africa. Only a few dispersed genomes originated from ^W^West-Africa.

To verify the geographic separation within L5 and L6, we conducted an independent phylogeographic analysis using the GenGIS software, where each whole genome SNP phylogeny was superimposed onto the five main African regions defined previously (Fig. 4A and C). We found several orientations of the tree’s geographical axis resulting in less crossings than expected by chance in L6 (p<0.001, 10.000 permutations; Fig. 4D). By contrast, for L5 we did not find any lineage axis with less crossing than expected by chance (Fig. 4B). These results indicate a marked geographical structure within L6 but not within L5. To further verify the different phylogeographical structures within L5 and L6, we calculated population differentiation indices considering each African region as a different population for each lineage. This analysis revealed some phylogeographical substructure within L6, where the percentage of variation attributed to different regions within Africa was 15% (Fst=0.15, p<0.00001). By contrast, L5 did not show any well-marked population differentiation, as the percentage of the variance attributed to population differences was only 6.6 %, with the rest of the variation attributed to intra-population differences (Fst=0.036, p<0.00001). This result further supports the observation of higher geographical structure within L6 than L5.

**Fig. 4.**
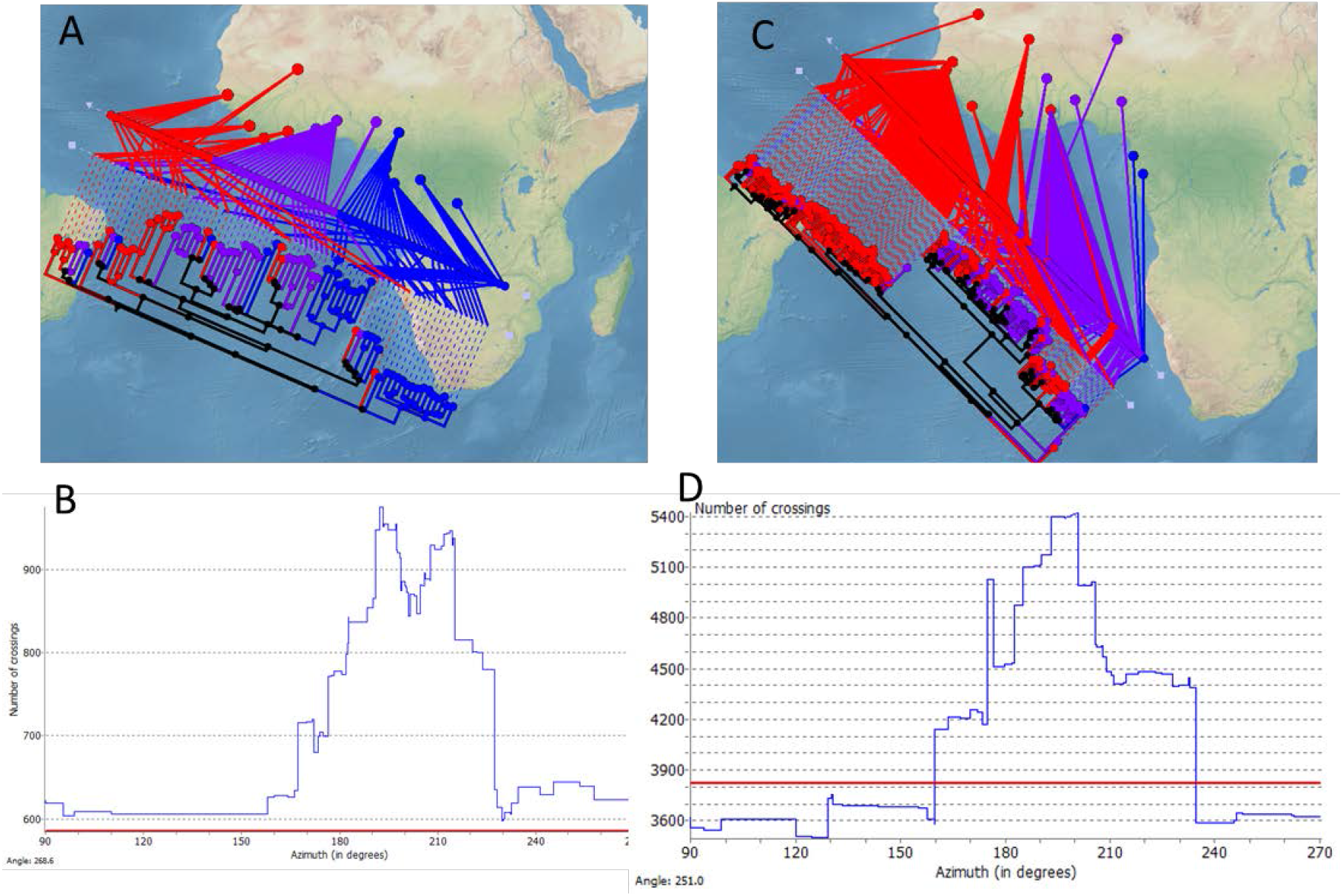
Phylogeographical structure in L5 and L6. Linear axis plot between the genomic phylogeny and the geographical origin of the genomes for L5 (A) and L6 (C). Histograms show the number of crossing for each inclination of the axis, and the red line indicates the number of crossing expected by chance for L5 (B) and L6 (D).

Finally, we explored possible differences in geographic range. Our dataset was geographically biased because it was designed to assemble as many L5 and L6 genomes from as many countries as possible. We therefore analysed our genome dataset together with two other large datasets where samples were not genome sequenced but genotyped using spoligotyping to compare the geographic distributions of L5 and L6 (*33*, *38*). This combined dataset included N=733 L5 from 27 African countries and N=1,031 L6 from 18 African countries. We expected that a broader geographical distribution of a specific lineage associated with a lower probability that two individuals selected randomly will belong to the same country. We used the Simpson’s Index (D) to measure the probability that two individuals randomly selected from a sample will belong to the same country. We found a larger diversity of countries of origin in L5 than in L6 (D=2.29 vs D=1.78, t-test α<0.05) indicating a broader geographic distribution of L5.

### The ancestral geographical distribution of L5, L6 and L9

Next, we explored the most likely geographical origin of L5 and L6 using four methods based on a Bayesian approach (*39*). The probabilities of ancestral distribution areas for the principal nodes were always congruent with at least two methods, but the results of the two other methods were either inconclusive or showed minor discrepancies (Fig. 5A and Fig. S2). For L5, two of the four methods inferred ^E^West Africa as the most likely origin (marginal probability was 1.0 using both Bayesian binary and S-DIVA), while the other two were inconclusive (marginal probabilities were ^E^West - Central: 0.94 and 0.58 with Bay Area and DEC, respectively; node 783 in Fig. 5A and Fig. S2). For L6, two methods also pointed to ^E^West Africa as the most likely origin (0.77, 1.0, of marginal probability using Bayesian binary and S-DIVA, respectively) and two methods supported both regions of West Africa as equally likely (0.94 and 0.58 using Bay area and DEC, respectively; node 592 in Fig. 5A and Fig. S2). The ancestral distribution of L9 was predicted to be East Africa based on all four methods (node 396 in Fig. 5A and Fig. S2).

**Fig. 5.**
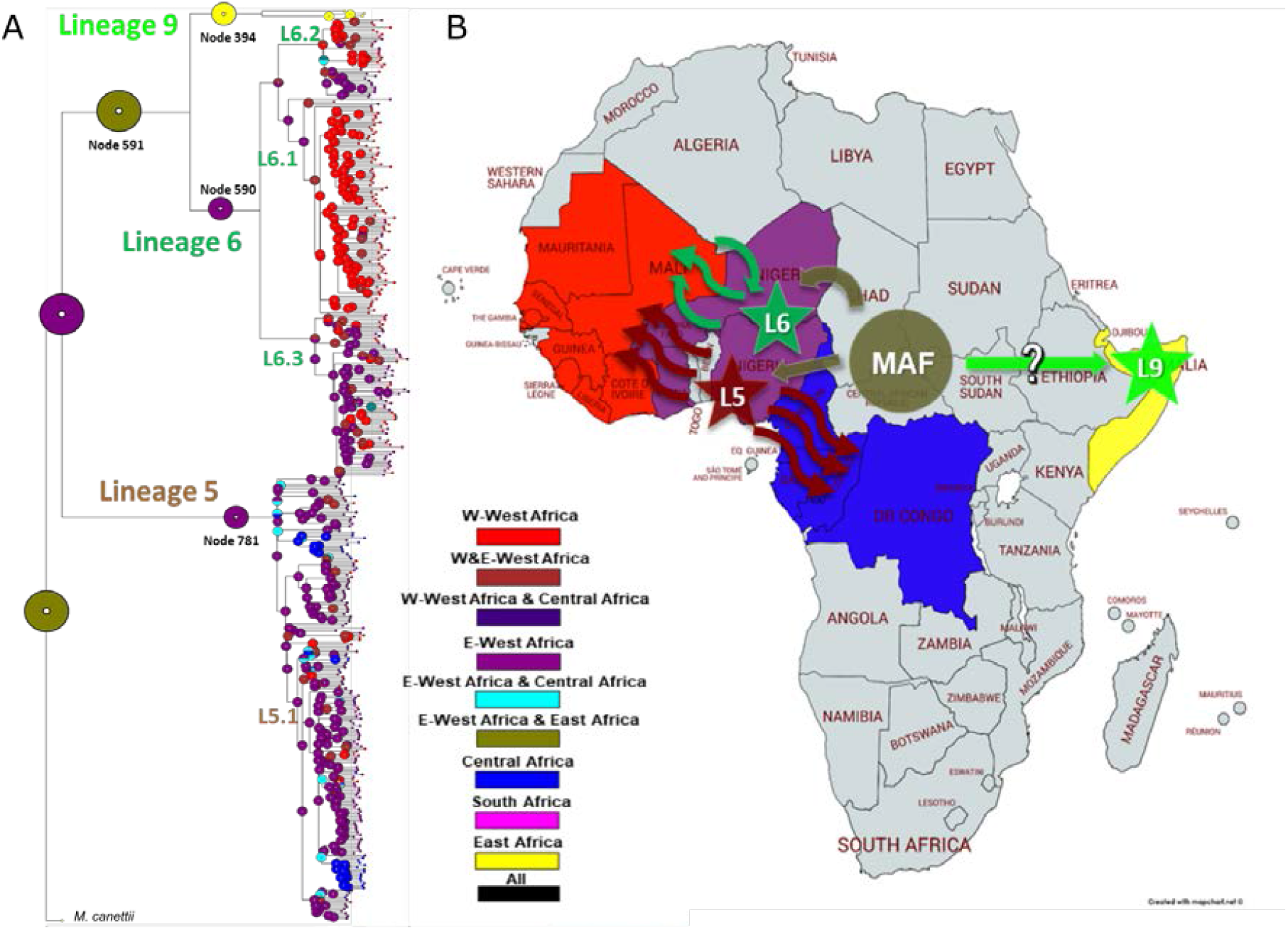
Geographical ancestral distributions of L5, L6 and L9. A. Ancestral area reconstruction by the Bayesian binary model onto the maximum likelihood phylogeny. Circles represent the probabilities of ancestral ranges, and the most likely ancestral areas are indicated by their corresponding colour code. B. Four geographical areas considered in this analysis are coloured in the map, the most likely areas ancestral areas for each lineage shown as stars, and movements of strains inferred from phylogeny indicated as arrows.

The ancestral distribution of the common ancestor between L6 and L9 was not that clearly predicted because of marginal probabilities of the methods supporting ^E^West Africa (0.65, 0.57 using BMBM, and DEC; node 591 Fig. 5A and Fig. S2A) and two methods supporting both regions in West Africa (0.5 using S-DIVA and Bay Area).

By contrast, the ancestral distribution for L5, L6 and L9 showed more consistency, where ^E^West Africa was supported by three methods (0.74, 1.0 and 0.57 using S-DIVA, BMBM and DEC, respectively) and one method predicting both ^E^West Africa and East Africa with a marginal probability of 0.99 (Bay Area: node 784 Fig. 5A and Fig. S4).

### Differences in genetic diversity between lineages

In support of our previous findings based on a more limited dataset (*40*), we found that L6 was significantly more genetically diverse than L5 with significantly higher number of SNPs between pairs of sequences (median values 553 vs 321; p-value < 2.2e-15), and significantly higher average nucleotide diversity (1.4×10-4 vs 8.7×10-5; p-value < 2.2e-15). To explore if this trend was consistent across the whole genome, we studied the nucleotide diversity in different regions that might be under different selection pressures: essential genes, non-essential genes, antigens, and T cell epitopes (Fig. 6). Although the genetic diversity was higher in all these different gene categories for L6 (Fig. 6), epitopes showed an inverted pattern in diversity between lineages (Fig. 6). Specifically, epitopes in L6 showed significantly higher genetic diversity than non-essential genes (Wilcoxon signed rank test p-value < 2.2e-15), while the opposite was found for L5, with epitopes showing significantly lower genetic diversity than non-essential genes (Wilcoxon signed rank test p-value < 2.2e-15).

**Fig. 6.**
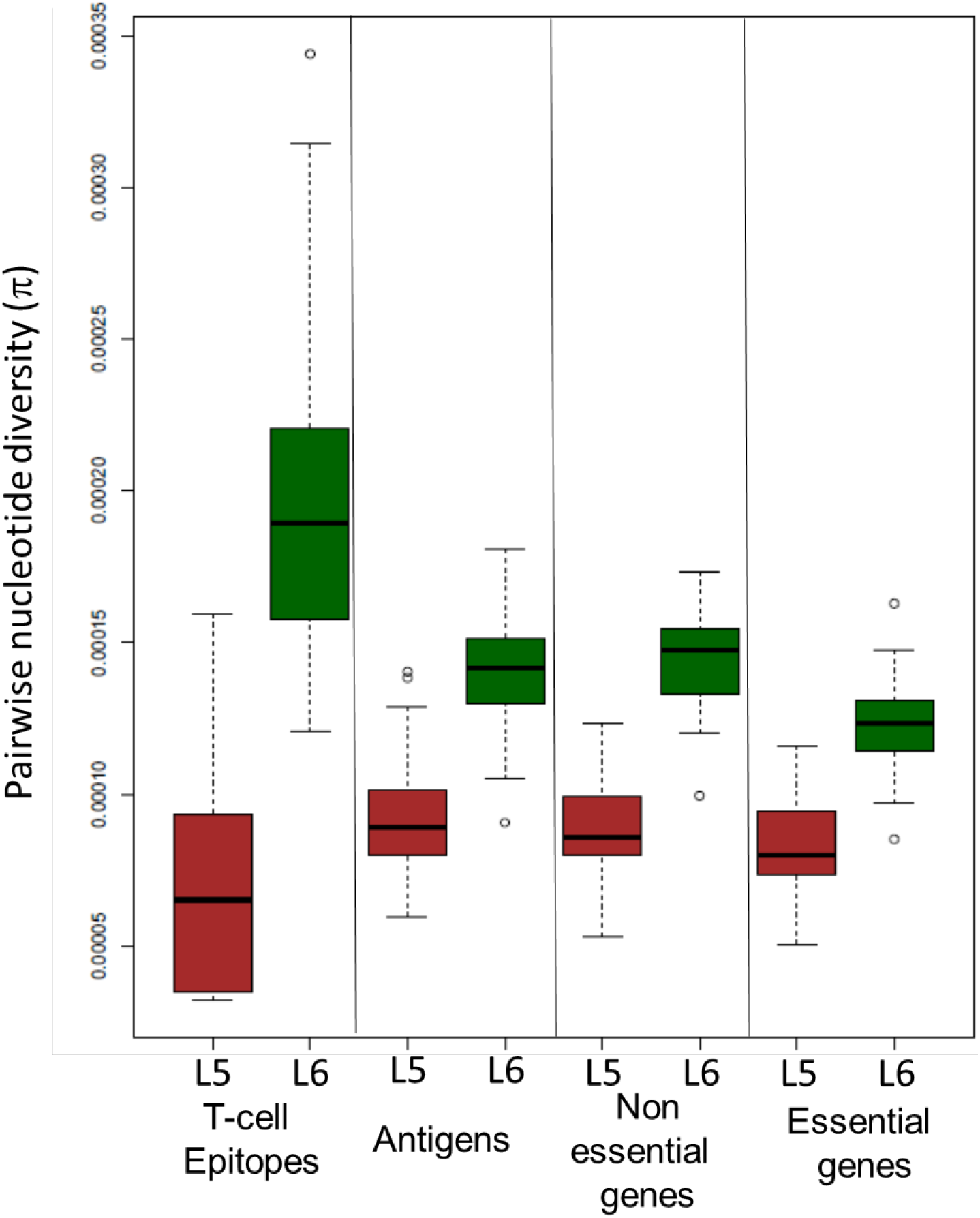
Nucleotide diversity (π). Comparison of pairwise nucleotide diversity (π) between L5 and L6 across gene categories.

### Drug resistance mutations

Antibiotic pressure is a strong selective force in bacteria including MTBC. Hence, we explored the difference in drug resistance determinants between L5 and L6. We found that among the 424 genomes analysed, 89 (21%) showed at least one genetic marker of antimycobacterial drug resistance, with 24 (6%) being multi-drug resistant (which is resistance to at least isoniazid and rifampicin, Table S8). The most common resistance found was for streptomycin, with 60 genomes showing 13 different resistance-conferring mutations. The next most common was resistance to rifampicin and isoniazid, with 32 and 29 genomes, respectively. Additional resistance was found to ethambutol, fluoroquinolones, ethionamide, pyrazinamide and aminoglycosides (Table S8). L5 genomes were more likely than L6 genomes to carry mutations associated with any resistance (OR 2.05 [95% confidence interval (CI) 1.26-3.31], p-value=2.29×10^−3^ using Fisher’s Exact Test). However, this was not due to a single antibiotic resistance profile because both lineages did not differ significantly when comparing the number of drug resistance mutations to fluoroquinolones (p-value=0.32), ethambutol (p-value=0.32), isoniazid (p-value=0.32), rifampicin (p-value=0.2), or streptomycin (p-value=0.34). Contrary to a previous report by Ates et al. (*36*), we found no evidence of differences in drug resistance genotype between L5.2 and other L5 genomes (OR 1.21 [95% CI 0.36-4.11], p-value=0.49, Fisher’s Exact Test).

## Discussion

*M. africanum* has traditionally been considered a single entity and a separate species from what classically has been referred to as *M. tuberculosis* sensu stricto. The results presented here provide novel insights into the genomic particularities of the different lineages within *M. africanum*: L5, L6 and a new group described in this study, L9. Differences between these three lineages further emphasize the need to consider these lineages as separate phylogenetic and ecologic variants within the MTBC.

Unexpectedly, our study of the global diversity of *M. africanum* revealed the presence of another MTBC lineage in Africa: L9, which is genetically close to L6. But unlike L5 and L6 that predominately occur in West Africa, L9 seems to be restricted to the East of Africa. Given that only five L9 isolates were included in our study, future studies are needed to confirm this observation (*7*, *8*). In this respect, L9 is similar to L7 and the recently described L8 (*3*), which are also mainly restricted to East Africa, but genetically more distant. We found clinical strains of L9 to be rare compared to L5 and L6, and this observation also resembles the situation for L7 and L8. As mentioned, we cannot dismiss the notion that this might be due to limited sampling, but the observation that clinical strains from L7, L8 and L9 originate in East Africa and are generally rare, while L5 and L6 are more prevalent and distributed across West and Central, raises the question of whether the reduced prevalence of L7, L8 and L9 is due to biological reasons, or social-environmental causes that renders L7, L8 and L9 to be less successful. The lack of experimental and epidemiological data on L7, L8 and L9 impedes a profound discussion on the matter. However, the fact that L9 is genetically closer to L6 and L5 than to L7 and L8, speaks against a common intrinsic biological determinant shared by L7, L8 and L9. Instead, convergence in the biology of the strains and/or in the socio-demography of the host is a more likely driver of the evolutionary history of L7, L8 and L9.

Our phylogeographic analyses localized the common ancestors of L5 and L6 to ^E^West Africa. We could detect that several subgroups of L5 moved from ^E^West Africa to Central Africa, while L6 subgroups moved mostly within West Africa. One of these events resulted in half of the L6 genomes in our dataset moving from ^E^West Africa to ^W^West Africa and with few dispersals back to ^E^West Africa (Fig. 5B). The ancestral reconstruction of L6 and L9 did not provide any clear signal, with ^E^West Africa and East Africa equally supported. For the ancestral distribution of all *M. africanum,* there was no consensus, but three out of four methods agreed on ^E^West Africa being the most likely place of origin. That would imply that L5 and L6 diversified there, and L9 migrated to East Africa. Remarkably, the fact that L9 is only present in East Africa, similar to L7 and L8 (*3*, *7*, *8*), suggests either one or two migration events to East Africa, depending on the ancestral distribution of the MTBC as a whole (Fig. 5B). Because *M. canettii*, the most closely related species of *M. tuberculosis* is highly restricted to East Africa, we and others have proposed that East Africa is the likely origin of the MTBC (*41*–*43*). If confirmed, the current geographical distribution of L5, L6 and L9 could be explained by a migration of their common ancestor from East Africa to West Africa, with the ancestor of L9 then moving back to East Africa. Alternatively, if the origin of the MTBC was in Central or West Africa, the current distribution would reflect at least three migration events to East Africa: one for the ancestor of L8, one for the ancestor of L7 and one for the ancestor of L9.

The work presented here also demonstrates differences in the population structure of L5 compared to L6. While L6 showed a marked phylogenetic structure comprising distinct sublineages associated with different geographical regions, the classification of L5 into sublineages was not so clearly supported despite the fact that L5 showed broader geographical range compared to L6. Additionally, our work confirms previous observations of differences in the genomic diversity, where L6 shows a higher diversity compared to L5 (*40*). In particular, human T cell epitopes in L6 were more diverse than non-essential genes, while the opposite was true for L5. Several studies have shown that human T cell epitopes in the human-adapted MTBC are overall more conserved than non-essential genes (*44*–*46*). This observation gave rise to the hypothesis that the MTBC might benefit from T cell recognition that drives lung pathology, leading to enhanced bacterial transmission (*47*). The fact that L6 differs in this respect from L5 and the other human-adapted MTBC lineages, indicates a potential different ecological niche, including possible animal reservoirs (*17*), which would also be supported by the phylogenetic proximity of L6 to the animal-adapted lineages of the MTBC (Figure 1).

We found L5 genomes more likely to carry any drug resistance-conferring mutations than L6. This result was consistent with previous findings from Ghana where L5 was compared to L4 (*48*). Due to the dominance of L5 genomes from Ghana in our dataset, we cannot rule out that our observation might have been partially driven by the Ghanaean genomes. However, unlike the previous report from Ghana, our study found L5 to be associated with any resistance, as opposed to specifically with a single antibiotic. In addition, contrary to the previous study from Ates et al. (*36*) based on a smaller dataset, our larger sampling indicated no association between drug resistance and a specific sublineage of L5 (*36*).

Our main study limitation is sampling bias, leading to an overrepresentation of isolates from the Gambia and Ghana. The overrepresentation of genomes from these two countries could contort our observation regarding the genomic diversity and population structure. Moreover, including more genomes from other countries will likely reveal additional sub-lineages within L5 and L6. Importantly, as stated in the previous paragraph, the association between L5 and drug resistance can be partially driven by a similar situation reported in Ghana previously. However, the differences we found between L5 and L6 is unlikely driven by this overrepresentation, because each country was enriched with strains from one of the two lineages.

In summary, we describe a large-scale whole-genome sequencing and a comprehensive phylogenomic analysis of clinical isolates classically referred to as “*M. africanum”* from 21 countries across Africa. Our findings have unravelled hidden diversity, a complex evolutionary history, and differential patterns of variation between lineages. Our results contribute to a better understanding of the MTBC lineages restricted to parts of Africa. These findings might assist in unraveling the molecular signatures of adaptations, and inform the development of targeted interventions for controlling TB in that part of the world.

## Methods

### *M. africanum* dataset

We analysed 697 L5 and L6 genomes to determine the genetic diversity, phylogeography and population structure of *M. africanum* (Table S1). This dataset included 495 newly sequenced genomes and 88 genomes from a previous study (*4*). Geographical origin was determined as the country of origin of the patient, and when not available the country of isolation. Because the number of different countries was too high to be shown properly in the figures, and some of them only included very few genomes, we grouped countries together into five African regions following definitions in (*38*): three big regions such as South, East and Central Africa, and two regions within West Africa, where most of the isolates come from. Western part of West Africa includes Gambia, Senegal, Mauritania, Sierra Leone, Liberia, Guinea, Ivory Coast, and Mali while the Eastern part of West Africa includes Ghana, Nigeria, Benin Niger, Burkina Faso). African maps were built using Mapcharnet^®^ (https://mapchart.net/africa.html)

### Bacterial Culture, DNA extraction and Whole-Genome sequencing

Archived MTBC isolates were revived by sub-culturing on Lowenstein Jensen media slants supplemented with 0.4% sodium pyruvate or with 0.3% glycerol to enhance the growth of the different lineages and incubated at 37 °C. Five loops full of colonies were harvested at the late exponential phase into 2 mL cryo-vials containing 1 mL of sterile nuclease-free water, inactivated at 98 °C for 60 minutes for DNA extraction using the previously described hybrid DNA extraction method (*48*). The MTBC lineages were then confirmed by spoligotyping and long sequence polymorphisms and sent for whole genome sequencing.

The MTBC isolates were grown in 7H9-Tween 0.05% medium (BD) +/− 40mM sodium pyruvate. We extracted genomic DNA after harvesting the bacterial cultures in the late exponential phase of growth using the CTAB method (*49*).

Sequencing libraries were prepared using NEXTERA XT DNA Preparation Kit (Illumina, San Diego, USA). Multiplexed libraries were paired-end sequenced on Illumina HiSeq2500 (Illumina, San Diego, USA) with 151 or 101 cycles when sequenced at the Genomics Facility Basel, HiSeq 2500 (100 bp, paired end) when sequenced at the Wellcome Sanger Institute, or on Illumina MiSeq (250 and 300 bp, paired end) or NextSeq (150 bp, paired end) according to the manufacturer’s instruction (Illumina, San Diego, USA) when sequenced at the genomics facilities in Research Center Borstel.

#### Bioinformatics analysis

##### Mapping and variant calling of Illumina reads

The obtained FASTQ files were processed with Trimmomatic v 0.33 (SLIDINGWINDOW: 5:20) (*50*) to clip Illumina adaptors and trim low quality reads. Any reads shorter than 20 bp were excluded for the downstream analysis. Overlapping paired-end reads were merged with SeqPrep v 1.2 (overlap size = 15) (https://github.com/jstjohn/SeqPrep). We used BWA v 0.7.13 (mem algorithm) (*51*) to align the resultant reads to the reconstructed ancestral sequence of *M. tuberculosis* obtained in (*44*). Duplicated reads were marked by the Mark Duplicates module of Picard v 2.9.1 (https://github.com/broadinstitute/picard) and excluded. To avoid false positive calls, Pysam v 0.9.0 (https://github.com/pysam-developers/pysam) was used to exclude reads with an alignment score lower than (0.93*read_length)-(read_length*4*0.07)), corresponding to more than 7 miss-matches per 100 bp. SNPs were called with Samtools v 1.2 mpileup (*52*) and VarScan v 2.4.1 (*53*) using the following thresholds: minimum mapping quality of 20, minimum base quality at a position of 20, minimum read depth at a position of 7X and without strand bias. Only SNPs considered to have reached fixation within an isolate were considered (at a within-host frequency of ≥90%). Conversely, when the within-isolate SNP frequency was ≤10% the ancestor state was called. Mixed infections or contaminations were discarded by excluding genomes with more than 1000 variable positions with within-host frequencies between 90% and 10% and genomes for which the number of within-host SNPs was higher than the number of fixed SNPs. Additionally, we excluded genomes with average read depth < 15 X (after all the referred filtering steps). All SNPs were annotated using snpEff v4.11 (*54*), in accordance with the *M. tuberculosis* H37Rv reference annotation (NC_000962.3). SNPs falling in regions such as PPE and PE-PGRS, phages, insertion sequences and in regions with at least 50 bp identities to other regions in the genome were excluded from the analysis as in (*55*). Customized scripts were used to calculate mean coverages per gene corrected by the size of the gene. Gene deletions were determined as regions with no coverage to the reference genome.

### Phylogenetic reconstruction and ancestry estimation

All 695 genomes were used to produce an alignment containing only polymorphic sites. The alignment was used to infer a maximum likelihood phylogenetic tree using the MPI parallel version of RAxML (*56*). We used the General Time Reversible model of nucleotide substitution under the Gamma model of rate heterogeneity and performed 1000 alternative runs on distinct starting trees combined with rapid bootstrap inference. To correct the likelihood for ascertainment bias introduced by only using polymorphic site, we used Lewis correction (*57*). The software Tremmer (*32*) was used to remove redundancy in the collection of 695 whole genome SNP alignment with the stop option - *RTL* 0.95, i.e. keeping 95% of the original tree length. The resulting reduced dataset of 424 genomes was kept for subsequent analysis. First we used the reduced dataset plus a collection of 35 representative animal genomes to produce an alignment containing only polymorphic sites and inferred a maximum likelihood phylogenetic tree as described above. The best-scoring Maximum Likelihood topology is shown. The phylogeny was rooted using *Mycobacterium canettii*. The topology was annotated and coloured using the package *ggtree* (*58*) from R (*59*) and InkScape.

We inferred the biogeographic histories of L5 and L6 using Statistical-Dispersal Analysis (S-DIVA) and Bayesian Binary MCMC (BBM) Method For Ancestral State, Dispersal-Extinction-Cladogenesis (DEC), and Bayesian inference for discrete Areas (BayArea) implemented in RASP v4.0 (*39*). Because we did not have the geographical origin of 18 samples, we used a phylogeny containing only samples from Africa where the isolation or place of birth of the patient was known. The possible ancestral ranges at each node on a selected tree were obtained. For S-DIVA the number of maximum areas was kept as 2. For BBM analysis, chains were run simultaneously for 500000 generations. The state was sampled every 100 generations. Estimated Felsenstein 1981 + Gamma was used with null root distribution.

### Population structure and genetic diversity

Genetic structure indices and corrected pairwise SNP differences between the five African regions where genomes are grouped (Western West Africa, Easter West Africa, Central Africa, South Africa, and East Africa) were calculated using Analysis of MOlecular VAriance (AMOVA) using information on the allelic content of haplotypes, as well as their frequencies implemented in Arlequin 3.5.2.2 (*60*). The significance of the covariance components was tested using 20000 permutations by non-parametric permutation procedures.

Pairwise SNP differences and mean nucleotide diversity per site (π) was calculated using the R package *ape* (*61*). π was calculated as the mean number of pairwise mismatches among L5 and L6 divided by the total length of queried genome base pairs, which comprise the total length of the genome after excluding repetitive regions (see above) (*62*). Confidence intervals for π were obtained by bootstrapping (1000 replicates) by re-sampling with replacement the nucleotide sites of the original alignments of polymorphic positions using the function *sample* in R (*59*). Lower and upper levels of confidence were obtained by calculating the 2.5th and the 97.5th quantiles of the π distribution obtained by bootstrapping. Population structure was evaluated using Principal Component Analysis (PCA) on SNP differences using *adegenet* (*63*) and plotted using the *plot* function in R.

To further explore geographical structure we evaluated the relation between the genomic phylogeny and the geographical origin of the genomes for each lineage separately using linear axis analysis in GenGISvs2.2.2 (*64*). The default GenGIS Africa map was used and a maximum likelihood phylogenetic tree was constructed from whole genome SNPs as described above for each lineage separately. A linear axis plot (10000 permutations) was run at significance level p-value = 0.001. If there is geographical separation, we expect the geographical distribution of the genomes to fit the phylogenetic tree structure. Fitting the tree is determined by finding a linear axis where the ordering of leaf nodes matches the ordering of sample sites according to the geographical distribution of each leaf node. If we draw a line between each leaf nodes in the phylogeny and its geographical distribution, a perfect match will result in minimum crossing of lines. Consequently, marked phylogeographical structure will result in significantly less crossing than the number of crossings expected by chance.

Simpson’s Index (D) for geographical diversity were calculated using three different datasets: 1) the current dataset (N=424), 2) 489 L5 and L6 strains obtained from the SITVIT2 database (*33*), a publicly available database that contains available genotyping (spoligotyping and MIRU-VNTRs), demographic and epidemiologic information on 111,635 clinical isolates, and 3) 837 genomes genotyped as L5 and L6 from 3580 strains from West Africa (*38*).

### Antimycobacterial resistance determining mutations and genes

We have used a compiled list of resistant mutations for 11 antibiotics compiled from two independent curated datasets (*65*).

## General

Library preparation and sequencing was done in the Genomics Facility at ETH Zürich, Basel, Switzerland. Calculations were performed at sciCORE (http://scicore.unibas.ch/) scientific computing core facility at University of Basel.

## Author contributions

Mireia Coscolla: Conceptualization, Data curation, Formal analysis, Investigation, Methodology, Visualization, Writing – original draft

Chloe Marie Loiseau: Data curation, Methodology, Writing -review & editing

Daniela Brites: Data curation, Conceptualization, Investigation, Formal Analysis, Writing -review & editing

Fabrizio Menardo Formal analysis, Investigation, Writing -review & editing

Sonia Borrell: Resources, Writing -review & editing

C. D’Nira Sanoussi: Investigation, Writing – review & editing

Conor Meehan: Conceptualization, Investigation, Writing – review & editing

Isaac Darko Otchere: Data curation, Formal analysis, Writing – review & editing

Leonor Sanchez-Busó: Data curation, Writing – review & editing

Julian Parkhill: Conceptualization, Supervision, Writing -review & editing

Patrick Beckert: Data curation, Writing – review & editing.

Stefan Niemann: Resources, Supervision, Writing -review & editing

Dissou Affolabi: Resources, Writing – review & editing

Prince Assare: Data curation, Formal analysis, Writing – review & editing

Florian Gehre: Data curation, review & editing, Writing – review & editing

Martin Antonio: Resources, Writing – review & editing

Adwoa Asante-Poku: Data curation, Writing – review & editing

Paula Ruíz-Rodriguez: Visualization, Methodology, Writing – review & editing

Janet Fyfe: Resources, Writing – review & editing

Robin Kobbe: Resources, Writing – review & editing

Martin P. Grobusch: Resources, Writing – review & editing

Abraham S. Alabi: Resources, Writing – review & editing

Lukas Fenner: Resources, Writing – review & editing

Erik C. Boettger: Resources, Writing – review & editing

Beisel Christian: Methodology, Writing – review & editing

Simon Harris: Conceptualization, Funding acquisition, Project administration, Supervision, Writing – review & editing

Dorothy Yeboah-Manu: Conceptualization, Funding acquisition, Project administration, Supervision, Writing – review & editing

Bouke de Jong: Conceptualization, Funding acquisition, Resources, Supervision, Project administration, Writing – review & editing,

Sebastien Gagneux: Conceptualization, Funding acquisition, Resources, Supervision, Project administration, Writing – original draft.

## Data and materials availability

All raw data generated for this study have been submitted to the European Genome-phenome Archive (EGA; https://www.ebi.ac.uk/ega/) under the study accession numbers PRJEB38317 and PRJEB38656. Individual runs accession number for new and published sequences are indicated in Table S1.

## Supplementary figures

**Fig. S1.**
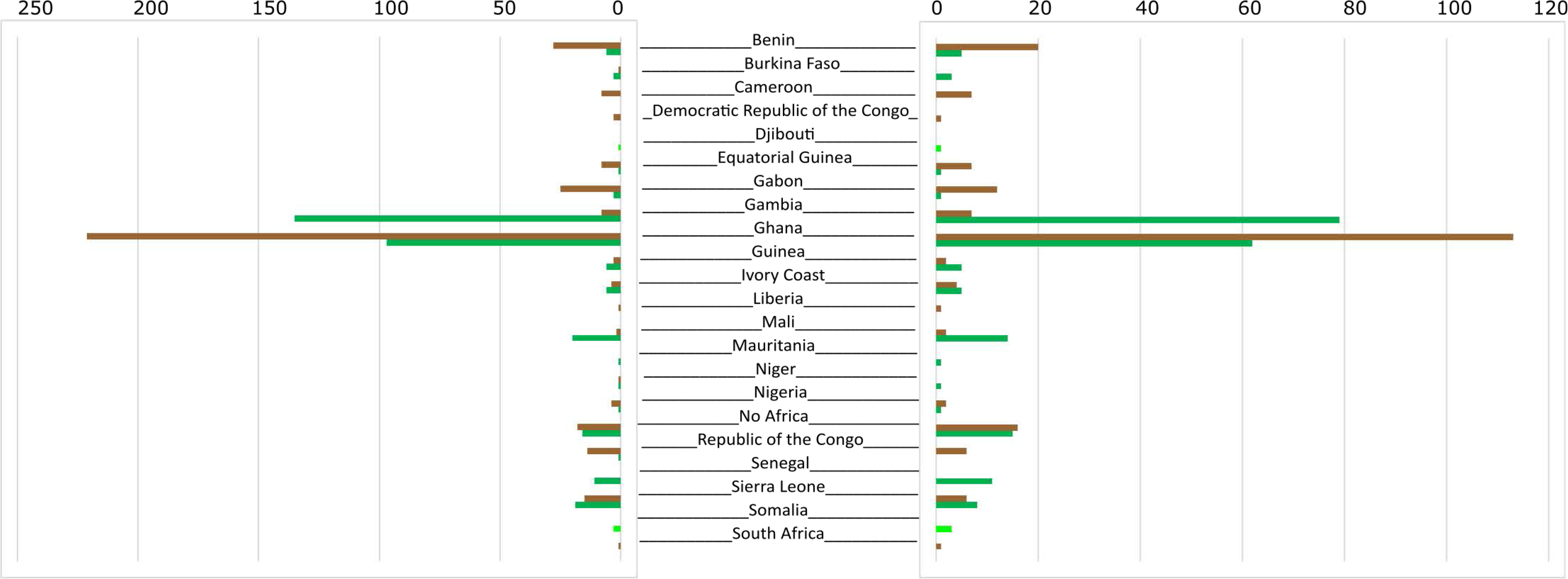
Lineage and country distribution. Genomes analysed for the initial dataset (A) and the non-redundant dataset (B). L5 genomes are indicated in brown bars, L6 genomes in green bars and L9 genomes in light green bars.

**Fig. S2.**
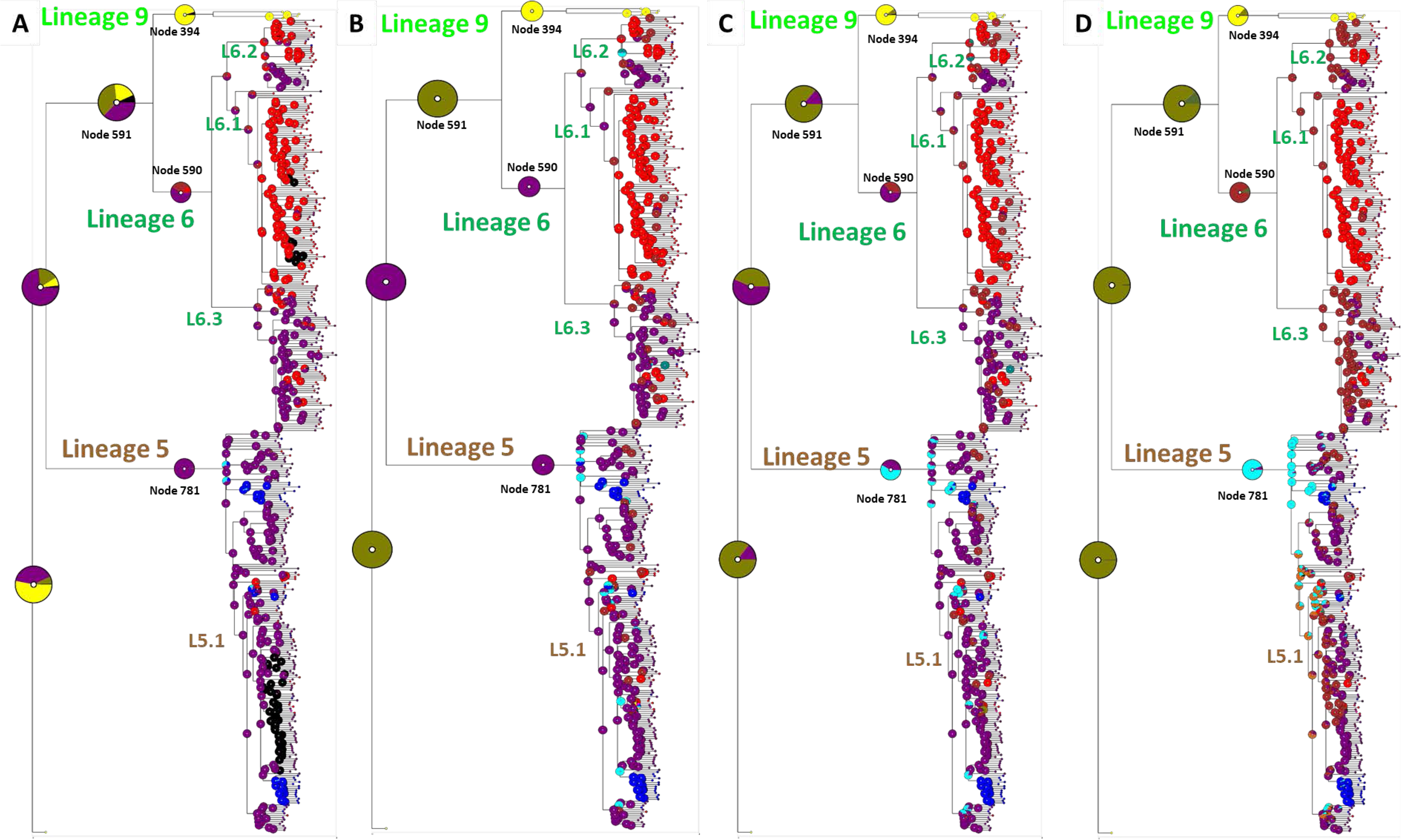
Ancestral area reconstruction onto the maximum likehood phylogeny. Circles represent the probabilities of ancestral ranges, and the most likely ancestral areas are indicated by their corresponding color code. The inset map represents the four geographical areas considered in this analysis. Results for all four methods are shown: Bayesian binary (A), DIVA (B) DEC (C) and BayArea (C).

## Supplementary tables

**Table S1. Genomes analysed.** Genome identifier, sequence accession numbers and sequencing statistics.

**Table S2. L9 specific mutations.** Synonymous and non-synonymous mutations in all lineage 9 genomes and absent in other strains from the dataset.

**Table S3. Pairwise F_ST_ values for L5 sublineages.**

**Table S4. Population average pairwise differences between L5 sublineages.**

**Table S5. Pairwise F_ST_ values for L6 sublineages.**

**Table S6. Population average pairwise differences between L6 sublineages.**

**Table S7. Sublineages SNPs.** SNPs defining L5 and L6 sublineages. SNPs in previously reported drug resistant genes were excluded.

**Table S8. Drug resistance mutations and genomes harbouring those mutations.**

## References

1. W. H. Organization, “Global tuberculosis report 2019” (ISBN 978-92-4-156571-4, 2019).

2. M. A. Riojas, K. J. McGough, C. J. Rider-Riojas, N. Rastogi, M. H. Hazbon, Phylogenomic analysis of the species of the *Mycobacterium tuberculosis* complex demonstrates that *Mycobacterium africanum, Mycobacterium bovis, Mycobacterium caprae, Mycobacterium microti* and *Mycobacterium pinnipedii* are later heterotypic synonyms of *Mycobacterium tuberculosis*. Int J Syst Evol Microbiol 68, 324–332 (2018).

3. J. C. Semuto Ngabonziza, C. Loiseau, M. Marceau, A. Jouet, F. Menardo, O. Tzfadia, R. Antoine, E. B. Niyigena, W. Mulders, K. Fissette, M. Diels, C. Gaudin, S. Duthoy, W. Ssengooba, E. André, M. K. Kaswa, Y. M. Habimana, D. Brites, D. Affolabi, J. B. Mazarati, B. C. de Jong, L. Rigouts, S. Gagneux, C. J. Meehan, P. Supply, A sister lineage of the *Mycobacterium tuberculosis* complex discovered in the AfricanGreat Lakes region. Nature communications Aceepted, (2020).

4. D. Brites, C. Loiseau, F. Menardo, S. Borrell, M. B. Boniotti, R. Warren, A. Dippenaar, S. D. C. Parsons, C. Beisel, M. A. Behr, J. A. Fyfe, M. Coscolla, S. Gagneux, A New Phylogenetic Framework for the Animal-Adapted *Mycobacterium tuberculosis* Complex. Front Microbiol 9, 2820 (2018).

5. S. Gagneux, Ecology and evolution of *Mycobacterium tuberculosis*. Nat Rev Microbiol 16, 202–213 (2018).

6. S. Gagneux, K. DeRiemer, T. Van, M. Kato-Maeda, B. C. de Jong, S. Narayanan, M. Nicol, S. Niemann, K. Kremer, M. C. Gutierrez, M. Hilty, P. C. Hopewell, P. M. Small, Variable host-pathogen compatibility in *Mycobacterium tuberculosis*. Proceedings of the National Academy of Sciences 103, 2869–2873 (2006).

7. R. Firdessa, S. Berg, E. Hailu, E. Schelling, B. Gumi, G. Erenso, E. Gadisa, T. Kiros, M. Habtamu, J. Hussein, J. Zinsstag, B. D. Robertson, G. Ameni, A. J. Lohan, B. Loftus, I. Comas, S. Gagneux, R. Tschopp, L. Yamuah, G. Hewinson, S. V. Gordon, D. B. Young, A. Aseffa, Mycobacterial Lineages Causing Pulmonary and Extrapulmonary Tuberculosis, Ethiopia. Emerging infectious diseases 19, 460–463 (2013).

8. Y. Blouin, Y. Hauck, C. Soler, M. Fabre, R. Vong, C. Dehan, G. Cazajous, P. L. Massoure, P. Kraemer, A. Jenkins, E. Garnotel, C. Pourcel, G. Vergnaud, Significance of the Identification in the Horn of Africa of an Exceptionally Deep Branching *Mycobacterium tuberculosis* Clade. PLoS ONE 7, (2012).

9. B. C. de Jong, M. Antonio, S. Gagneux, *Mycobacterium africanum*-Review of an Important Cause of Human Tuberculosis in West Africa. Plos Neglected Tropical Diseases 4, (2010).

10. W. H. Haas, G. Bretzel, B. Amthor, K. Schilke, G. Krommes, S. Rusch-Gerdes, V. Sticht-Groh, H. J. Bremer, Comparison of DNA fingerprint patterns of isolates of *Mycobacterium africanum* from east and west Africa. J Clin Microbiol 35, 663–666 (1997).

11. M. Kato-Maeda, P. J. Bifani, B. N. Kreiswirth, P. M. Small, The nature and consequence of genetic variability within *Mycobacterium tuberculosis*. The Journal of Clinical Investigation 107, 533–537 (2001).

12. B. Ofori-Anyinam, A. J. Riley, T. Jobarteh, E. Gitteh, B. Sarr, T. I. Faal-Jawara, L. Rigouts, M. Senghore, A. Kehinde, N. Onyejepu, M. Antonio, B. C. de Jong, F. Gehre, C. J. Meehan, Comparative genomics shows differences in the electron transport and carbon metabolic pathways of *Mycobacterium africanum* relative to *Mycobacterium tuberculosis* and suggests an adaptation to low oxygen tension. Tuberculosis (Edinb) 120, 101899 (2020).

13. M. Coscolla, S. Gagneux, Consequences of genomic diversity in *Mycobacterium tuberculosis*. Seminars in Immunology 26, 431–444 (2014).

14. P. Asare, A. Asante-Poku, D. A. Prah, S. Borrell, S. Osei-Wusu, I. D. Otchere, A. Forson, G. Adjapong, K. A. Koram, S. Gagneux, D. Yeboah-Manu, Reduced transmission of *Mycobacterium africanum* compared to *Mycobacterium tuberculosis* in urban West Africa. International journal of infectious diseases: IJID: official publication of the International Society for Infectious Diseases 73, 30–42 (2018).

15. M. Huet, N. Rist, G. Boube, D. Potier, [Bacteriological study of tuberculosis in Cameroon]. Rev Tuberc Pneumol (Paris) 35, 413–426 (1971).

16. G. Kallenius, T. Koivula, S. Ghebremichael, S. E. Hoffner, R. Norberg, E. Svensson, F. Dias, B. I. Marklund, S. B. Svenson, Evolution and clonal traits of *Mycobacterium tuberculosis* complex in Guinea-Bissau. J Clin Microbiol 37, 3872–3878 (1999).

17. B. C. de Jong, M. Antonio, S. Gagneux, *Mycobacterium africanum*—Review of an Important Cause of Human Tuberculosis in West Africa. PLoS Negl Trop Dis 4, (2010).

18. S. Homolka, E. Post, B. Oberhauser, A. G. George, L. Westman, F. Dafae, S. Rüsch-Gerdes, S. Niemann, High genetic diversity among *Mycobacterium tuberculosis* complex strains from Sierra Leone. BMC Microbiol 8, 103 (2008).

19. B. C. de Jong, I. Adetifa, B. Walther, P. C. Hill, M. Antonio, M. Ota, R. A. Adegbola, Differences between tuberculosis cases infected with *Mycobacterium africanum*, West African type 2, relative to Euro-American *Mycobacterium tuberculosis*: an update. FEMS Immunology & Medical Microbiology 58, 102–105 (2010).

20. A. Asante-Poku, D. Yeboah-Manu, I. D. Otchere, S. Y. Aboagye, D. Stucki, J. Hattendorf, S. Borrell, J. Feldmann, E. Danso, S. Gagneux, *Mycobacterium africanum* Is Associated with Patient Ethnicity in Ghana. PLoS Negl Trop Dis 9, e3370 (2015).

21. A. Asante-Poku, I. D. Otchere, S. Osei-Wusu, E. Sarpong, A. Baddoo, A. Forson, C. Laryea, S. Borrell, F. Bonsu, J. Hattendorf, C. Ahorlu, K. A. Koram, S. Gagneux, D. Yeboah-Manu, Molecular epidemiology of *Mycobacterium africanum* in Ghana. BMC Infect Dis 16, 385 (2016).

22. D. Brites, S. Gagneux, Co-evolution of *Mycobacterium tuberculosis* and Homo sapiens. Immunological Reviews 264, 6–24 (2015).

23. C. G. Meyer, G. Scarisbrick, S. Niemann, E. N. Browne, M. A. Chinbuah, J. Gyapong, I. Osei, E. Owusu-Dabo, T. Kubica, S. Rusch-Gerdes, T. Thye, R. D. Horstmann, Pulmonary tuberculosis: virulence of *Mycobacterium africanum* and relevance in HIV co-infection. Tuberculosis (Edinb) 88, 482–489 (2008).

24. B. Diarra, M. Kone, A. C. G. Togo, Y. D. S. Sarro, A. B. Cisse, A. Somboro, B. Degoga, M. Tolofoudie, B. Kone, M. Sanogo, B. Baya, O. Kodio, M. Maiga, M. Belson, S. Orsega, M. Krit, S. Dao, Maiga, II, R. L. Murphy, L. Rigouts, S. Doumbia, S. Diallo, B. C. de Jong, *Mycobacterium africanum* (Lineage 6) shows slower sputum smear conversion on tuberculosis treatment than *Mycobacterium tuberculosis* (Lineage 4) in Bamako, Mali. PLoS One 13, e0208603 (2018).

25. M. C. Hlavsa, P. K. Moonan, L. S. Cowan, T. R. Navin, J. S. Kammerer, G. P. Morlock, J. T. Crawford, P. A. LoBue, Human Tuberculosis due to Mycobacterium bovis in the United States, 1995–2005. Clinical Infectious Diseases 47, 168–175 (2008).

26. D. Park, H. Qin, S. Jain, M. Preziosi, J. J. Minuto, W. C. Mathews, K. S. Moser, C. A. Benson, Tuberculosis due to Mycobacterium bovis in Patients Coinfected with Human Immunodeficiency Virus. Clinical Infectious Diseases 51, 1343–1346 (2010).

27. B. C. de Jong, P. C. Hill, R. H. Brookes, J. K. Otu, K. L. Peterson, P. M. Small, A. Adegbola, “*Mycobacterium africanum*: a new opportunistic pathogen in HIV infection?” in Aids (England, 2005), vol. 19, pp. 1714–1715.

28. N. D. C. Sanoussi, B. C. deJong, M. Odoun, K. Arekpa, M. Ali Ligali, O. Bodi, S. Harris, B. Ofori-Anyinam, D. Yeboah-Manu, I. Otchere, Darko, A. Asante-Poku, Gagneux, M. Coscolla, L. Rigouts, D. Affolbi, Low sensitivity of the MPT64 identification test to detect lineage 5 of the *Mycobacterium tuberculosis* complex. J Med Microbiol 67, 1718–1727 (2018).

29. B. C. de Jong, P. C. Hill, R. H. Brookes, S. Gagneux, D. J. Jeffries, J. K. Otu, S. A. Donkor, A. Fox, K. P. McAdam, P. M. Small, R. A. Adegbola, *Mycobacterium africanum* elicits an attenuated T cell response to early secreted antigenic target, 6 kDa, in patients with tuberculosis and their household contacts. J Infect Dis 193, 1279–1286 (2006).

30. B. Ofori-Anyinam, F. Kanuteh, S. C. Agbla, I. Adetifa, C. Okoi, G. Dolganov, G. Schoolnik, O. Secka, M. Antonio, B. C. de Jong, F. Gehre, Impact of the Mycobaterium africanum West Africa 2 Lineage on TB Diagnostics in West Africa: Decreased Sensitivity of Rapid Identification Tests in The Gambia. PLOS Neglected Tropical Diseases 10, e0004801 (2016).

31. S. Lipworth, R. Jajou, A. de Neeling, P. Bradley, W. van der Hoek, G. Maphalala, M. Bonnet, E. Sanchez-Padilla, R. Diel, S. Niemann, Z. Iqbal, G. Smith, T. Peto, D. Crook, T. Walker, D. van Soolingen, SNP-IT Tool for Identifying Subspecies and Associated Lineages of *Mycobacterium tuberculosis* Complex. Emerg Infect Dis 25, 482–488 (2019).

32. F. Menardo, C. Loiseau, D. Brites, M. Coscolla, S. M. Gygli, L. K. Rutaihwa, A. Trauner, C. Beisel, S. Borrell, S. Gagneux, Treemmer: a tool to reduce large phylogenetic datasets with minimal loss of diversity. BMC Bioinformatics 19, 164 (2018).

33. D. Couvin, A. David, T. Zozio, N. Rastogi, Macro-geographical specificities of the prevailing tuberculosis epidemic as seen through SITVIT2, an updated version of the *Mycobacterium tuberculosis* genotyping database. Infection, Genetics and Evolution, (2018).

34. R. Brosch, S. V. Gordon, M. Marmiesse, P. Brodin, C. Buchrieser, K. Eiglmeier, T. Garnier, C. Gutierrez, G. Hewinson, K. Kremer, L. M. Parsons, A. S. Pym, S. Samper, D. van Soolingen, S. T. Cole, A new evolutionary scenario for the *Mycobacterium tuberculosis* complex. Proc Natl Acad Sci U S A 99, 3684–3689 (2002).

35. S. Mostowy, D. Cousins, J. Brinkman, A. Aranaz, M. A. Behr, Genomic deletions suggest a phylogeny for the *Mycobacterium tuberculosis* complex. J Infect Dis 186, 74–80 (2002).

36. L. S. Ates, A. Dippenaar, F. Sayes, A. Pawlik, C. Bouchier, L. Ma, R. M. Warren, W. Sougakoff, L. Majlessi, J. W. J. van Heijst, F. Brossier, R. Brosch, Unexpected Genomic and Phenotypic Diversity of *Mycobacterium africanum* Lineage 5 Affects Drug Resistance, Protein Secretion, and Immunogenicity. Genome biology and evolution 10, 1858–1874 (2018).

37. S. Wright, Genetical Structure of Populations. Nature 166, 247–249 (1950).

38. F. Gehre, S. Kumar, L. Kendall, M. Ejo, O. Secka, B. Ofori-Anyinam, E. Abatih, M. Antonio, D. Berkvens, B. C. de Jong, A Mycobacterial Perspective on Tuberculosis in West Africa: Significant Geographical Variation of *M. africanum* and Other *M. tuberculosis* Complex Lineages. PLoS Negl Trop Dis 10, e0004408 (2016).

39. Y. Yu, A. J. Harris, C. Blair, X. He, RASP (Reconstruct Ancestral State in Phylogenies): a tool for historical biogeography. Mol Phylogenet Evol 87, 46–49 (2015).

40. I. D. Otchere, M. Coscolla, L. Sanchez-Buso, A. Asante-Poku, D. Brites, C. Loiseau, C. Meehan, S. Osei-Wusu, A. Forson, C. Laryea, A. I. Yahayah, A. Baddoo, G. A. Ansa, S. Y. Aboagye, P. Asare, S. Borrell, F. Gehre, P. Beckert, T. A. Kohl, S. N’Dira, C. Beisel, M. Antonio, S. Niemann, B. C. de Jong, J. Parkhill, S. R. Harris, S. Gagneux, D. Yeboah-Manu, Comparative genomics of *Mycobacterium africanum* Lineage 5 and Lineage 6 from Ghana suggests distinct ecological niches. Sci Rep 8, 11269 (2018).

41. R. Hershberg, M. Lipatov, P. M. Small, H. Sheffer, S. Niemann, S. Homolka, J. C. Roach, K. Kremer, D. A. Petrov, M. W. Feldman, S. Gagneux, High Functional Diversity in *Mycobacterium tuberculosis* Driven by Genetic Drift and Human Demography. PLoS Biology 6, e311 (2008).

42. P. Supply, M. Marceau, S. Mangenot, D. Roche, C. Rouanet, V. Khanna, L. Majlessi, A. Criscuolo, J. Tap, A. Pawlik, L. Fiette, M. Orgeur, M. Fabre, C. Parmentier, W. Frigui, R. Simeone, E. C. Boritsch, A. S. Debrie, E. Willery, D. Walker, M. A. Quail, L. Ma, C. Bouchier, G. Salvignol, F. Sayes, A. Cascioferro, T. Seemann, V. Barbe, C. Locht, M. C. Gutierrez, C. Leclerc, S. D. Bentley, T. P. Stinear, S. Brisse, C. Medigue, J. Parkhill, S. Cruveiller, R. Brosch, Genomic analysis of smooth tubercle bacilli provides insights into ancestry and pathoadaptation of *Mycobacterium tuberculosis*. Nature Genetics 45, 172–179 (2013).

43. I. Comas, M. Coscolla, T. Luo, S. Borrell, K. E. Holt, M. Kato-Maeda, J. Parkhill, B. Malla, S. Berg, G. Thwaites, D. Yeboah-Manu, G. Bothamley, J. Mei, L. Wei, S. Bentley, S. R. Harris, S. Niemann, R. Diel, A. Aseffa, Q. Gao, D. Young, S. Gagneux, Out-of-Africa migration and Neolithic coexpansion of *Mycobacterium tuberculosis* with modern humans. Nat Genet 45, 1176–1182 (2013).

44. I. Comas, J. Chakravartti, P. M. Small, J. Galagan, S. Niemann, K. Kremer, J. D. Ernst, S. Gagneux, Human T cell epitopes of *Mycobacterium tuberculosis* are evolutionarily hyperconserved. Nat Genet 42, 498–503 (2010).

45. M. Coscolla, R. Copin, J. Sutherland, F. Gehre, B. de Jong, O. Owolabi, G. Mbayo, F. Giardina, J. D. Ernst, S. Gagneux, *M. tuberculosis* T Cell Epitope Analysis Reveals Paucity of Antigenic Variation and Identifies Rare Variable TB Antigens. Cell Host Microbe 18, 538–548 (2015).

46. C. S. Lindestam Arlehamn, S. Paul, F. Mele, C. Huang, J. A. Greenbaum, R. Vita, J. Sidney, B. Peters, F. Sallusto, A. Sette, Immunological consequences of intragenus conservation of *Mycobacterium tuberculosis* T-cell epitopes. Proceedings of the National Academy of Sciences of the United States of America 112, E147–E155 (2015).

47. J. D. Ernst, The immunological life cycle of tuberculosis. Nat Rev Immunol 12, 581–591 (2012).

48. I. D. Otchere, A. Asante-Poku, S. Osei-Wusu, A. Baddoo, E. Sarpong, A. H. Ganiyu, S. Y. Aboagye, A. Forson, F. Bonsu, A. I. Yahayah, K. Koram, S. Gagneux, D. Yeboah-Manu, Detection and characterization of drug-resistant conferring genes in *Mycobacterium tuberculosis* complex strains: A prospective study in two distant regions of Ghana. Tuberculosis (Edinb) 99, 147–154 (2016).

49. J. T. Belisle, M. G. Sonnenberg, Isolation of genomic DNA from mycobacteria. Methods Mol Biol 101, 31–44 (1998).

50. A. M. Bolger, M. Lohse, B. Usadel, Trimmomatic: a flexible trimmer for Illumina sequence data. Bioinformatics 30, 2114–2120 (2014).

51. H. Li, R. Durbin, Fast and accurate long-read alignment with Burrows-Wheeler transform. Bioinformatics 26, 589–595 (2010).

52. H. Li, A statistical framework for SNP calling, mutation discovery, association mapping and population genetical parameter estimation from sequencing data. Bioinformatics 27, 2987–2993 (2011).

53. D. C. Koboldt, Q. Zhang, D. E. Larson, D. Shen, M. D. McLellan, L. Lin, C. A. Miller, E. R. Mardis, L. Ding, R. K. Wilson, VarScan 2: somatic mutation and copy number alteration discovery in cancer by exome sequencing. Genome Res 22, 568–576 (2012).

54. P. Cingolani, A. Platts, L. Wang le, M. Coon, T. Nguyen, L. Wang, S. J. Land, X. Lu, D. M. Ruden, A program for annotating and predicting the effects of single nucleotide polymorphisms, SnpEff: SNPs in the genome of Drosophila melanogaster strain w1118; iso-2; iso-3. Fly (Austin) 6, 80–92 (2012).

55. D. Stucki, D. Brites, L. Jeljeli, M. Coscolla, Q. Liu, A. Trauner, L. Fenner, L. Rutaihwa, S. Borrell, T. Luo, Q. Gao, M. Kato-Maeda, M. Ballif, M. Egger, R. Macedo, H. Mardassi, M. Moreno, G. T. Vilanova, J. Fyfe, M. Globan, J. Thomas, F. Jamieson, J. L. Guthrie, A. Asante-Poku, D. Yeboah-Manu, E. Wampande, W. Ssengooba, M. Joloba, W. H. Boom, I. Basu, J. Bower, M. Saraiva, S. E. Vasconcellos, P. Suffys, A. Koch, R. Wilkinson, L. Gail-Bekker, B. Malla, S. D. Ley, H. P. Beck, B. C. de Jong, K. Toit, E. Sanchez-Padilla, M. Bonnet, A. Gil-Brusola, M. Frank, V. N. Penlap Beng, K. Eisenach, I. Alani, P. W. Ndung’u, G. Revathi, F. Gehre, S. Akter, F. Ntoumi, L. Stewart-Isherwood, N. E. Ntinginya, A. Rachow, M. Hoelscher, D. M. Cirillo, G. Skenders, S. Hoffner, D. Bakonyte, P. Stakenas, R. Diel, V. Crudu, O. Moldovan, S. Al-Hajoj, L. Otero, F. Barletta, E. J. Carter, L. Diero, P. Supply, I. Comas, S. Niemann, S. Gagneux, *Mycobacterium tuberculosis* lineage 4 comprises globally distributed and geographically restricted sublineages. Nat Genet 48, 1535–1543 (2016).

56. A. Stamatakis, RAxML-VI-HPC: maximum likelihood-based phylogenetic analyses with thousands of taxa and mixed models. Bioinformatics 22, 2688–2690 (2006).

57. P. O. Lewis, A likelihood approach to estimating phylogeny from discrete morphological character data. Syst Biol 50, 913–925 (2001).

58. Y.Guangchuang, S. D. K., Z. Huachen, G. Yi, L. T. Tsan-Yuk, ggtree: an r package for visualization and annotation of phylogenetic trees with their covariates and other associated data. Methods in Ecology and Evolution 8, 28–36 (2017).

59. R. C. Team. (Vienna, Austria, 2018).

60. L. Excoffier, G. Laval, S. Scheneider, Arlequin (version 3.0): An integrated software package for population genetics data analysis. Evolutionary Bioinformatics 1, 47–50 (2005).

61. E. Paradis, J. Claude, K. Strimmer, APE: Analyses of Phylogenetics and Evolution in R language. Bioinformatics 20, 289–290 (2004).

62. D. L. Hartl, A. G. Clarck, Principles of population genetics (Sinauer Associates, Inc, Sunderland, MA, 2006).

63. T. Jombart, adegenet: a R package for the multivariate analysis of genetic markers. Bioinformatics 24, 1403–1405 (2008).

64. D. H. Parks, T. Mankowski, S. Zangooei, M. S. Porter, D. G. Armanini, D. J. Baird, M. G. Langille, R. G. Beiko, GenGIS 2: geospatial analysis of traditional and genetic biodiversity, with new gradient algorithms and an extensible plugin framework. PLoS One 8, e69885 (2013).

65. J. L. Payne, F. Menardo, A. Trauner, S. Borrell, S. M. Gygli, C. Loiseau, S. Gagneux, A. R. Hall, Transition bias influences the evolution of antibiotic resistance in *Mycobacterium tuberculosis*. PLoS Biol 17, e3000265 (2019).

